# SAXS/MC studies of the mixed-folded protein Cdt1 reveal monomeric, folded over conformations

**DOI:** 10.1101/2024.01.03.573975

**Authors:** Kyle P. Smith, Srinivas Chakravarthy, Amit Rahi, Manas Chakraborty, Kristen M. Vosberg, Marco Tonelli, Maximilian G. Plach, Arabela A. Grigorescu, Joseph E. Curtis, Dileep Varma

## Abstract

Cdt1 is a protein critical for DNA replication licensing and is well-established to be a binding partner of the minichromosome maintenance (MCM) complex. Cdt1 has also been demonstrated to have an emerging, “moonlighting” role at the kinetochore via direct binding to microtubules and to the Ndc80 complex. However, it is not known how the structure and conformations of Cdt1 could allow for these multiple, completely unique sets of protein complexes. And while there exist multiple robust methods to study entirely folded or entirely unfolded proteins, structure-function studies of combined, mixed folded/disordered proteins remain challenging. It this work, we employ multiple orthogonal biophysical and computational techniques to provide a detailed structural characterization of human Cdt1 92-546. DSF and DSCD show both folded winged helix (WH) domains of Cdt1 are relatively unstable. CD and NMR show the N-terminal and the linker regions are intrinsically disordered. Using DLS and SEC-MALS, we show that Cdt1 is polydisperse, monomeric at high concentrations, and without any apparent inter-molecular self-association. SEC-SAXS of the monomer in solution enabled computational modeling of the protein *in silico*. Using the program SASSIE, we performed rigid body Monte Carlo simulations to generate a conformational ensemble. Using experimental SAXS data, we filtered for conformations which did and did not fit our data. We observe that neither fully extended nor extremely compact Cdt1 conformations are consistent with our SAXS data. The best fit models have the N-terminal and linker regions extended into solution and the two folded domains close to each other in apparent “folded over” conformations. The best fit Cdt1 conformations are consistent with a function as a scaffold protein which may be sterically blocked without the presence of binding partners. Our studies also provide a template for combining experimental and computational biophysical techniques to study mixed-folded proteins.

## Introduction

Human Cdt1 is a 546-amino acid protein with two well-folded winged helix (WH) domains and two uncharacterized regions. These well-folded domains are referred to as the “middle” WH domain (WHM) and the C-terminal WH domain (WHC) (**Figure 1A**). Cdt1 was first discovered for its role in origin licensing during DNA replication, where it has been demonstrated to be critical for loading the MiniChromosome Maintenance (MCM) helicase onto the DNA, which in turn is required for DNA unwinding during replication. Even though the WH fold has been established to bind DNA in other contexts, intriguingly, the WHM and WHC domains of Cdt1 have not been demonstrated to directly bind DNA. It is still a part of the larger complex including the DNA-binding, Origin Recognition Complex (ORC), Cdc6, and MCM (the preRC) at the origins of replication on polymeric DNA (1,2). Interestingly, recent studies have shown that Cdt1 also performs an essential, “moonlighting” function during mitosis. It localizes to kinetochores of condensed chromosomes and to the microtubules of the mitotic spindle. This is thought to facilitate specific interactions between these two large mitotic complexes. Cdt1 seems to have retained certain aspects of its mitotic functionality, as it also functions in linking the kinetochore bound Ndc80 complex to polymeric microtubules, with other proteins in the assembly unit yet to be identified (3,4). Cdt1 has been enigmatic in that it does not possess any enzymatic activity. Recently, it has also been shown that the well-folded WHD’s of Cdt1 are crucial for its ability to bind microtubules, similar to that of another microtubule-binding protein complex, Ska1 (5,6). Interestingly, the Ska1 complex, like Cdt1, also associates with Ndc80 at kinetochores. It is unclear how the structure of Cdt1 can mediate specific, defined interactions with a multitude of other proteins, and underscore the importance of investigating its structural properties, which is currently missing. Further, the alteration of Cdt1 has been critically linked to genomic instability and developmental abnormalities such as primordial dwarfism syndrome (2,7). Given this, structure-function studies of Cdt1 could eventually lead to a better understanding of the pathology of these disease states.

**Figure 1.**
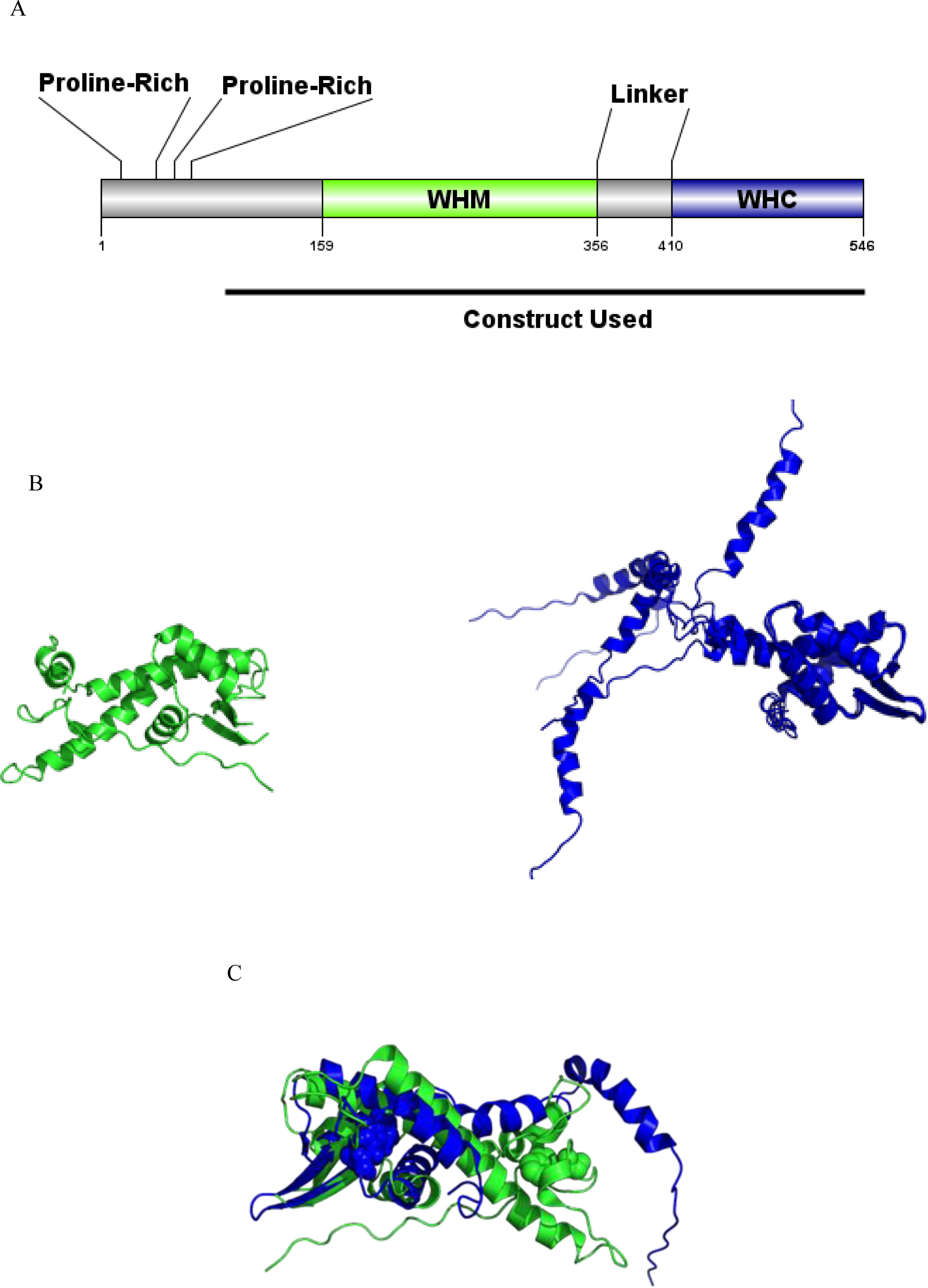
Domain map and published Cdt1 structures. (A) Full length protein and constructs employed in this and other published studies. (B) Crystal and NMR structures of the WHM (green) and WHC (blue) are shown as cartoons. (C) Overlay of WHM and WHC domains. Single tryptophan residue in each domain is shown in spheres.

The crystal structure of the mouse WHM domain was first determined in complex with its re-licensing inhibitor, geminin (8). Later, the structure of the WHM/geminin complex was also determined under higher-order, hetero-oligomeric states (8,9). Based on crystallography and SAXS data, the authors propose these different inhibitory complexes could allow for what they call “licensing-competent” and “licensing-defective states” for the WHM domain. Following the original WHM crystal structure, both the crystal and NMR structures of mouse Cdt1 WHC domain have also been published (10,11), but only as individual domains (**Figure 1B**). More recently, advances in single-particle Cryo Electron Microscopy (CryoEM) have allowed for a better understanding of Cdt1’s function in coordination with the MCM and ORC complexes as a part of the replication licensing machinery (12–15). However, notably missing from previous structural characterization of Cdt1 have been reductionist approaches to using solely the recombinant, multi-domain protein in isolation. While these existing cryoEM structures show one of the folded domains of Cdt1 in complex with the preRC, there is limited homology between yeast and human Cdt1. This data does not provide enough resolution for the structure of Cdt1 alone, an understanding of which is critical to elucidate how Cdt1 enters into complex formation with different multimeric protein assemblies during both DNA replication and mitosis. Notably, solution studies, as opposed to cryogenic ones, have been very limited.

As opposed to most folded proteins comprised of α-helices and β-sheets, other proteins exist as so-called intrinsically disordered proteins (IDP), which are generally flexible in solution. They do not form “classical” secondary structural elements and their structures were historically referred to as “random coil”. But while IDP’s are not well-folded, they also do not necessarily have completely random coil structure or are completely unfolded either and can have different local and global structures (16,17). It is estimated that approximately 50% of proteins in the human proteome include at least partial regions of intrinsically disorder (18,19). This estimate therefore includes proteins which contain both folded and unfolded portions, most commonly called “mixed-folded” proteins. The addition of folded domains almost certainly affects the structure of the disordered regions as well. For Cdt1, unsurprisingly, an amino acid sequence alignment generated by PRALINE shows the N-terminal and linker regions are less conserved than the folded WH domains (**Figure S1A_i_-A_iii_**, (20)). Significant work has supported the notion that disordered domains can bind to multiple partners with different structures (21). This, coupled with the accessibility and frequency of PTM’s, can enable mixed-folded proteins to function as scaffold proteins for signaling cascades and as protein-protein interaction “hubs” (22,23). Historically, disordered proteins were thought to be “undruggable”, but there have been some more recent cases in which small molecules can bind to, and alter the function of, disordered protein regions (24,25). And as a class, mixed-folded proteins have been implicated in schizophrenia (DISC1 (26)), polyglutamine disorders (Ataxin (27)), and microcephaly (Knl1 (28)), where treatment options are extremely limited.

Because mixed-folded proteins by definition have both well-folded and disordered regions, unlike structures generated from crystallography, NMR spectroscopy, and CryoEM, the structure of these proteins often cannot be captured using more static, higher-resolution techniques. And while machine learning approaches like AlphaFOLD (29), among others, have dramatically changed the structural understanding of folded proteins, they are not suitable for systems like Cdt1. Small Angle X-Ray Scattering (SAXS), however, is an in-solution scattering technique which has been applied to help characterize higher-order protein-protein complexes, protein folding, and IDP’s (30). Building upon this, molecular simulation via Molecular Dynamics (MD) and Monte Carlo (MC) approaches to model mixed-folded proteins have provided additional insight into the conformations of mixed-folded proteins. For example, in the case of HIV gag protein, SAXS/MD/MC approaches demonstrated the protein exists in a more compact conformation in solution, in contrast to an extended conformation required for simultaneous membrane and nucleic acid binding during virus assembly (31,32). Similarly, in the MCM complex, MD/MC approaches allowed for structure/function studies of the disordered regions and relative conformations of the folded domains (33,34). These historically difficult-to-study mixed-folded proteins must often be characterized by using complementary, orthogonal biophysical techniques to fully understand their structure/function. In this study, we use integrated biophysical, structural, and computational approaches, to describe the solution structural ensemble of the intrinsically mixed-folded protein Cdt1. Our modeled conformations suggest Cdt1 is folded over onto itself, with both WHM and WHC domains in close proximity to each other without any apparent inter-molecular self-interaction. Center-of-mass analysis validates these conformations. Our use of orthogonal, complementary biophysical and structural techniques also provides a template to study other mixed-folded protein systems.

## Materials and Methods

Certain commercial equipment, instruments, materials, suppliers, or software are identified in this paper to foster understanding. Such identification does not imply recommendation or endorsement by the National Institute of Standards and Technology, nor does it imply that the materials or equipment identified are necessarily the best available for the purpose. Unless otherwise indicated, all experiments were performed at the Northwestern University Department of Cell & Developmental Biology (Chicago, IL).

### Cloning and Molecular Biology

All His_6_-GFP-Cdt1 constructs and vectors used were generated as previously described (5). For protein expression, the constructs were transformed into BL21-CodonPlus (DE3)-RP competent cells (Agilent). For transformation and precultures, cells were grown in LB media. For unlabeled protein expression, proteins were grown in TPM media (20 g tryptone, 15 g yeast extract, 8 g sodium chloride, 2 g dibasic sodium phosphate, 1 g monobasic potassium phosphate per 1 L).

### Protein Expression and Purification

Conditions were optimized based on the initial protocols from Agarwal et al. (5). Several changes were made to increase yields. For protein expression, expression and purity were greatly reduced without a fresh BL21 transformation. The induction time was lowered to 3.5 hours to reduce proteolysis. After elution from Co2+ beads, the protein was concentrated to approximately 500 µL using centrifugal spin filters (Milipore) and injected over a Superdex 200 (S200) Increase 10/300 column (GE Healthcare) connected to an AKTA Pure FPLC (GE Healthcare). Under a flow rate of 0.5 mL/min, His_6_-GFP-Cdt1 92-546 eluted at approximately 11.5 mL. The primary contaminant was cleaved GFP. Approximately fractions 10.5-13.5 mL were pooled, flash frozen in liquid nitrogen, and stored at −80°C until use. ^15^N-labeled Cdt1 was purified identically to that of unlabeled protein. Protein was approximately 95% pure by SDS-PAGE (**Figure S1B**).

### Commercially Available Proteins

*B.taurus* carbonic anhydrase isozyme II from erythrocytes, *G.gallus* egg albumin (ovalbumin), *B.taurus* serine albumin, *S.cerevisiae* alcohol dehydrogenase, *O.cuniculus* muscle aldolase, *I.batatas* beta amylase, and *L.mesenteroides* blue dextran were purchased as lyophilized powders or preformulated solutions from Sigma-Aldrich and used without further purification.

### Protein Tag Cleavage and Buffer Equilibration

For His_6_-GFP tag cleavage, 800-1400 µL of approximately 5 mg/mL of protein was incubated with 100 µL of 2 mg/mL TEV protease, provided by the Northwestern Recombinant Protein Production Core. Each reaction was performed in a 1.5 mL tube. The tubes were gently rotated for 1 hour at room temperature (approximately 22 °C). Samples were put on ice, concentrated to approximately 300 µL and injected over a Superdex 200 Increase (S200) 10/300 column. Using a flow rate of 0.5 mL/min, cleaved Cdt1 92-546 eluted at approximately 12.2 mL and 11.5-12.5 mL were pooled, concentrated, and flash frozen in liquid nitrogen. To buffer exchange the proteins into the appropriate assay buffer before assay use, SEC using an S200 was employed to both buffer exchange and ensure folded, active samples distinct from potential aggregates. Protein concentration of tag-cleaved Cdt1 92-546 was calculated using the theoretical extinction coefficient (35) of all tryptophan, tyrosine, and reduced cysteine residues in our construct at 280 nm (21,930 M^−1^ cm^−1^) on a NanoDrop 1000 spectrophotometer (ThermoFischer).

### Cdt1 Expression for Isotopic Labeling

For ^15^N labeling of Cdt1, 1 L cultures of autoinduction media (Cold Spring Harbor Protocols) were used with ^15^N labeled ammonium chloride (Cambridge Isotope Laboratories). The final antibiotic concentration was increased to 0.1 mg/mL of kanamycin to reduce plasmid loss and poor yields. Cultures were grown in 2.5 L Ultra Yield™ flasks (Thomson Instrument Company) for increased aeration. 1 L cultures were inoculated with 5 mL of overnight preculture cells. The cultures were grown in darkness at 37 °C with shaking at 250 rpm until an OD_600_ of 0.6, when the temperature was lowered to 30 °C overnight. The cells continued to grow for approximately 16 additional hours. Cell pelleting, lysis and purification were identical to that of the unlabeled protein.

### TIRF Microscopy

A microscopy perfusion chamber was prepared by attaching an acid washed coverslip (22 × 22 mm, thickness 1 1/2, Corning) over a precleaned glass slide (75 × 25 mm, thickness 1 mm, Corning) using a double-sided tape (Scotch) resulting in a chamber volume of ∼10 µl. For more details, please see TIRF section in Afreen et al., 2022 [39]. Acid washed coverslips were further cleaned by holding them under flame for a couple of seconds prior to use. Solutions were then exchanged into the perfusion chamber by micropipette and filter paper. The chamber was activated by flowing in the following solution in an order in which they appear below, followed by incubation for 5 min and intermittent washing with 3 chamber volumes of buffer B (MRB80 supplemented with 10 µM taxol) : PLL PEG biotin (0.1 mg ml^−1^, Susos, AG), Streptavidin (0.625 mg ml^−1^, #S4762, Sigma), diluted solution of Taxol stabilized microtubules (1 in 100 dilutions form the stock), and κ-Casein (1 mg ml^−1^). All incubations were done inside a wet chamber to prevent evaporation. A solution of cleaved Cdt1 (100 nM, in buffer B) was subsequently flown into the chamber and incubated for 5 min. The chamber was then washed with the following antibody solutions anti Cdt1 (1 in 500 dilution in buffer B), and Alexa488 labeled-secondary antibody (1 in 1000 dilution in buffer B) with intermittent washing with buffer B. The chamber was finally washed with imaging buffer (MRB80 supplemented with 10 µM Taxol, 0.6 mg ml^−1^ κ-casein, 4 mM DTT, 50 mM glucose (#G8270, Sigma), 0.2 mg ml^−1^, catalase (#C9322, Sigma), and 0.4 mg ml^−1^glucose oxidase (#G7141, Sigma). Images in were recorded with the following microscopy settings. Imaging was performed with a Nikon Ti-2 inverted microscope equipped with Andor iXon3 CCD camera (Cambridge Scientific, Watertown, MA, USA), 1.49× NA, and 100× oil objective. The microscope produced a 512 × 512-pixel images with 0.16 µm per pixel resolution in both *x* and *y* directions. NIS-elements software (Nikon) was used for data acquisition with 100 ms exposure times. Microtubule images were collected using 640 nm excitation laser and images of Alexa 488 lagges antibody were collected using 488 nm laser excitation line.

### SEC-MALS

Experiments were performed at the Northwestern University Keck Biophysics Facility (Evanston, IL). Solution size exclusion chromatography coupled with multi-angle laser light scattering/quasi-elastic light scattering (SEC-MALS/QELS) experiments were conducted using Agilent Technologies 1200 LC HPLS system (Agilent Technologies, Santa Clara, CA) equipped with a Wyatt Dawn® Heleos™II 18-angle MALS light scattering detectors, Optilab® T-rEX™ refractive index detector, WyattQELS™ quasi-elastic (dynamic) light scattering (QELS) detector and ASTRA software (Wyatt Technology Europe GmbH). QELS interval was set to 0.5 sec and collection interval was set to 0.4 sec. Cdt1 92-546 was at a concentration of 3.53 mg/mL and was cleaved/purified the prior day before overnight storage at 4 °C. A total of 250 µL of each protein was injected and run on a Superdex 200 Increase (S200) 10/300 column (GE Healthcare) at a flow rate of 0.40 mL/min in SEC-MALS buffer (10 mM Tris pH 8.3, 150 mM NaCl, 0.25 mM TCEP) at room temperature (24.3°C). Monomeric bovine serum albumin was used to experimentally determine the dn/dc value (0.161 mL/g) under our conditions. The refractive index was set to 1.335 and the viscosity was set to 8.9450×10-3 g*cm-1*sec-1. R(h) analysis was based on Tayyab et. al. (36). Elution volume reported is at maximum A280 value. Void volume (Vo) was calculated using a 250 µL injection of 7.89 mg/mL Blue Dextran (Sigma-Aldrich). Inclusion volume (Vi) was calculated using a 250 µL injection of 2% ethanol (v/v). R(h) values were reported as previously published. Each protein standard was resuspended in SEC-MALS buffer to 2.0-8.0 mg/mL and syringe filtered through a 0.22 µm filter before injection of 250 µL.

### Intrinsic Fluorescence DSF (nanoDSF)

Experiments were performed at 2bind GmbH (Regensburg, Germany). Cdt1 and ovalbumin were diluted from stock concentration (6.35 mg/mL and 10 mg/mL, respectively) to 5 µM in “Stability Buffer” (25 mM HEPES pH 7.4, 150 mM NaCl). Duplicate samples were loaded into standard nanoDSF glass capillaries and analyzed on a Prometheus NT.48 device, equipped with additional back-reflection optics (NanoTemper Technologies, Munich, Germany). The fluorescence at 350 and 330 nm was measured while the temperature was increased from 20 to 95 °C at a rate of 1°C/min. Excitation power was set to 70%. Thermally induced aggregation (T_agg_) was also inferred by measuring light-scattering while increasing the temperature from 20 to 95°C at a rate of 1°C/min. Data were analyzed using PR. ThermControl (v2.0.4) software (NanoTemper Technologies, Munich, Germany). Thermal melting temperatures (T_m_) was inferred in first approximation from inflection points of 350/330 nm fluorescence ratio curves.

### CD Spectroscopy

Experiments were performed at the Northwestern University Keck Biophysics Facility (Evanston, IL). Data was collected on a Jasco J-815 CD Spectrophotometer (Jasco Analytical Instruments, Easton, MD). A flow rate of 60 ft^3^/hr nitrogen was used to purge the system of oxygen and was kept running throughout the experiments. Proteins were kept on ice before CD experiments to ensure they remained properly folded. For secondary structure assignment, approximately 75 µL of 1.16 mg/mL (54 µM) protein in CD Buffer (10 mM Tris pH 8.3, 150 mM NaCl, 0.5 mM TCEP) was put into a 0.1 mm slide cuvette (Hellma Analytics). Data integration time was set to 4 seconds, bandwidth was set to 2.0 mm, and 3 accumulations in “step” scanning mode. HT was also monitored and remained less than 600 V at all wavelengths. CD was recorded from 180 to 260 nm using a step size of 1 nm. For differential scanning measurements, 6 initial, preliminary spectra were collected to optimize path length, protein concentration, and buffer concentration. Final data collection was performed with 300 µl of 0.193 mg/mL (9 µM) protein in diluted CD Buffer (6 mM Tris pH 8.3, 90 mM NaCl, 0.3 mM TCEP) in a 1 mm cuvette (Hellma Analytics). The sample was heated from 10 °C to 90 °C with a 1°C/min temperature increase. The data were collected every 1°C at wavelengths of 209 and 220 nm, which corresponded local minima observed for the spectrum collected at 25°C. Temperature was held constant during data collection measurements. Data integration time was set to 1 second, bandwidth was set to 2.0 mm, and 1 accumulation in “continuous” scanning mode. HT was also monitored and remained less than 600 V at all wavelengths. CD was recorded from 195 to 280 nm. Data analysis was performed in the Spectra Manager® Ver.2 software package using the CD Pro Analysis and Protein Denaturation Analysis portions. CONTIN (37) was used to assign the secondary structure. Three CONTIN soluble protein databases [SP29 (178-260 nm), SP37 (185-240 nm) and SP43 (190-240 nm)] were used to fit the CD spectrum. The pecentages of secondary structures were averaged from fittings of each database. The root mean square deviation (RMSD) and normalized root mean square deviation (NMRSD) shows goodness of the fit and were around 3% and 2%, respectively. The melting temperatures extracted from the fit are within less than 0.5 °C difference (209 nm: 42.0° ± 0.2 and 220 nm: 42.4° ± 0.2).

### DLS

DLS measurements were performed using a Punk 0037 Dynamic Light Scattering Analyzer (UnChained Labs). Each sample was loaded into a disposable 5 µL sample holder (BladeCell) and measured at 20 °C. Intensity-weighted size distribution was obtained over ten runs made up of five measurements each. The solvent was recorded as PBS and its refractive index of 1.3 was assumed. The optical attenuator and laser intensity parameters were set automatically for each run. The reported R(h) was calculated from the average of three technical replicates using one protein production. One replicate was discarded because it had a poorer fit to the correlation function. One other replicate was discarded because it was 1.5x the standard deviation of the average of the other 3 replicates.

### NMR Spectroscopy

Experiments were performed at NMRFAM at the University of Wisconsin (Madison Wisconsin). ^15^N labeled Cdt1 92-546 was concentrated to 200 µM in 10 mM HEPES, 150 mM NaCl pH 7.4 buffer before dilution in 10% D2O (Sigma). ^15^N-HSQC data was collected on a 750 MHz Avance III HD spectrometer (Bruker) in Shigemi tubes (Shigemi).

### SEC-SAXS

Experiments were performed at the Advanced Photon Source BioCAT beamline (Sector 18-ID) of Argonne National Laboratory (Argonne, IL), as described by Mathew et al. (38). The 12-keV X-ray beam (λ = 1.033 Å) was focused on a 1.5-mm quartz capillary sample cell with 10µm walls. The scattering, in the momentum transfer range, *q* = 0.0038–0.4 Å^−1^, was collected on a Dectris Pilatus 3X 1M detector approximately 1.5 m downstream of the sample position. All instrumentation and experiments were performed at 25 °C. 275 μl of 5.64 mg/mL Cdt1 92-546 in storage buffer (25mM Tris pH 8.3, 500mM NaCl, 0.5mM TCEP, 5% sucrose (w/v)) was fed into the X-ray beam after passing through a Superdex 200 (S200) Increase 10/300 GL column (GE Healthcare) in SAXS buffer (50mM HEPES pH 7.4, 500mM NaCl, 1mM TCEP). A flow rate of 0.75 mL/min was used. The delay between protein emerging from the SEC column and its arrival at the beam position was approximately 1 minute. SAXS exposures with a length of 0.5 sec were collected every 3.0 sec. Scattering data was radially averaged, processed, scaled, and analyzed using PrimusQT (39). Exposures before and after sample elution were averaged and used as buffer background. Pair distribution functions, *P*(*R*), of the scattering centers were computed from the scattering curves using GNOM (40). Guiner analysis was performed using RAW (41).

### Molecular Simulations

The starting model was generated using the x-ray structure of human Cdt1 WHM (8), and from a human homology model of mouse Cdt1 (10). Residues 90-166 and 354-410 were added as initially linear sequences using PSFGEN and NAMD (42). The model was prepared using the CHARMM-32 force field (43). The initial structure was energy minimized and relaxed via 1 ns NVT molecular dynamics simulation. Subsequent Monte Carlo simulations were completed using the Monomer Monte Carlo module in the program SASSIE (41), with residues 90-163 and 351-443 denoted as “flexible”. 13724 out of an attempted 250,000 non-overlapping accepted structures were each energy minimized and used for subsequent analysis using SASSIE modules Chi-Square Filter and Density Plot. Analysis of domain distributions from the ensemble was done by calculating the center-of-mass for all atoms in the WHM (residues 160-350) and WHC (residues 410-546) domains for each structure in the trajectory.

### Graphics

Sequence alignment was visualized using PRALINE (20). Domain maps were drawn using DOG (44). Protein ribbon diagrams were visualized using PyMOL (45). Thermal stability, spectroscopy, SEC-MALS, and SEC-SAXS figures were visualized using Excel (Microsoft). Density plots were generated using SASSIE and visualized using VMD (46).

## Results and Discussion

### Human Cdt1 92-546 is competent in binding to microtubules

We previously predicted (5) the N-terminal 1-92 region of Cdt1 to be disordered partially based on its 18.5% proline content, which is approximately three times higher than across the human proteome (47). Therefore, all our experiments were conducted with the largest possible fragment (92–546), hereafter referred to just as “Cdt1” (**Figure 1A**). As a control protein, we chose to use ovalbumin, which has a similar molecular weight to Cdt1, is very well-folded with both alpha helices and beta sheets, and is very well-characterized structurally. We had previously demonstrated that a GFP-tagged version of Cdt1 92-546 binds directly to microtubules by using both microtubule copelleting as well as TIRF binding assays (5). To ensure that the cleaved Cdt1 92-546 molecule also retained the ability to bind microtubules, we carried out a Total Internal Reflection Fluorescence (TIRF) microscopy-based assay to test this. We observed that this tagless Cdt1 protein is able to bind to polymerized microtubules (**Figure 2A**). The methodology resembled an indirect immunostaining experiment (see Methods) where, we used a primary antibody to detect Cdt1 and then used a fluorescently labeled secondary antibody to visualize Cdt1 molecules that are bound to immobilized microtubules. This demonstrates our construct binds directly to microtubules, as expected.

**Figure 2.**
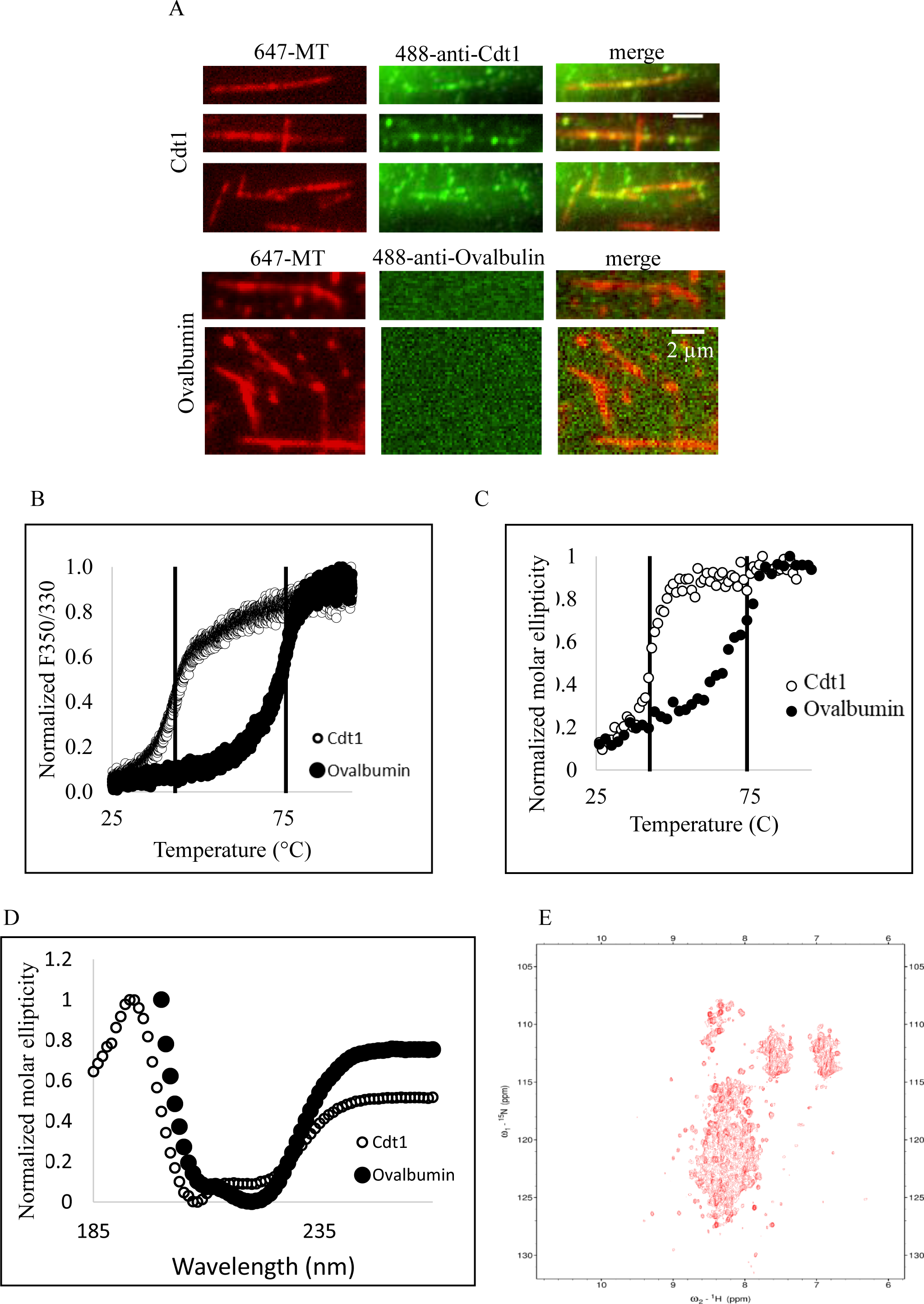
Microtubule binding, thermal denaturation and spectroscopic analyses of Cdt1. (A) TIRF analysis of Cdt1-microtubule binding. Scale bars represent 2 µm. (B) DSF of Cdt1 and ovalbumin. Normalized F350/330 ratio is plotted as a function of temperature. Vertical lines represent melting temperatures. Ovalbumin is plotted in closed circles and Cdt1 is plotted in open circles. (C) DSCD of Cdt1 and ovalbumin. Normalized molar ellipticity is plotted as a function of temperature. (D) CD spectra of Cdt1 and ovalbumin. Normalized CD is plotted as a function of wavelength. (E) ^15^N-^1^H TROSY HSQC of Cdt1. ^15^N and ^1^H signals are plotted in ppm.

### Thermal denaturation experiments demonstrate Cdt1 is unstable

To study the thermal stability of Cdt1, we performed differential scanning denaturation experiments. We used both circular dichroism (CD) and intrinsic fluorescence (Ex280, Em330/350) to measure the changes in secondary and tertiary elements, respectively. Our thermal stability data for ovalbumin (**Figure 2B** & **2C**) was similar to previously published data (48) given the difference in buffers (**Table 1** & **2**). For both DSF and DSCD, Cdt1 followed a simple two-state unfolding process. In fluorescence detection, we observed a melting temperature (T_m_) of 43 °C (**Figure 2B**). In CD, we calculated a T_m_ of 42°C (**Figure 2C**). Because each WH domain contains one buried tryptophan residue (**Figure 1C**) and has similar secondary structures, our denaturation data also supports the interpretation that the two domains have similar, low thermal stabilities. Given how low the Tm values are relative to 37°C, this demonstrates Cdt1 is quite intrinsically unstable. There are reports which anecdotally correlate low stability *in vitro* with low stability *in vivo* (49–51). It has been shown that Cdt1 is rapidly degraded at the end of G1 phase and that this is important in preventing genome reduplication (1,2). This low Tm *in vitro* could promote faster unfolding and proteosome degradation, which would have implications for Cdt1’s degradation during different stages of the cell cycle (52,53).

**Table 1:**
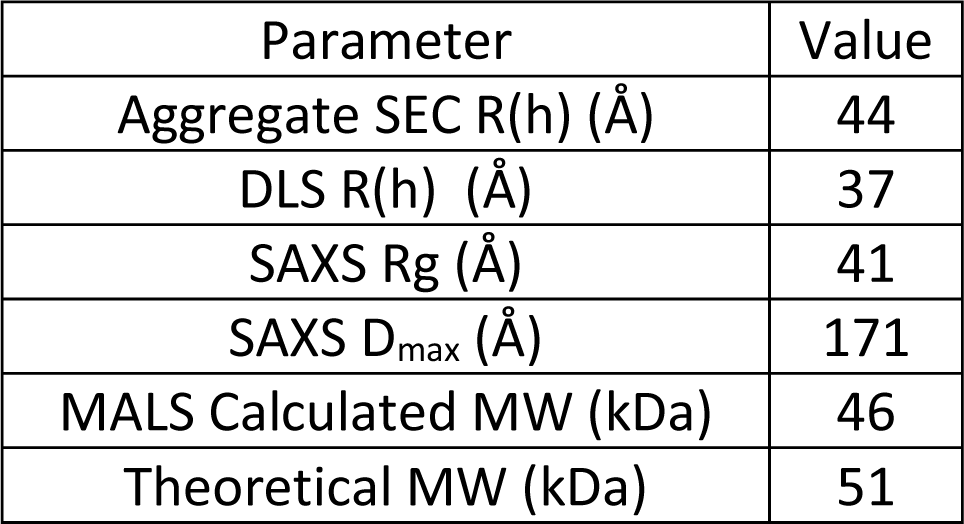
Biophysical properties of Cdt1.

| Parameter | Value |
| --- | --- |
| Aggregate SEC R(h) (Å) | 44 |
| DLS R(h) (Å) | 37 |
| SAXS R<sub>g</sub> (Å) | 41 |
| SAXS D<sub>max</sub> (Å) | 171 |
| MALS Calculated MW (kDa) | 46 |
| Theoretical MW (kDa) | 51 |

### Spectroscopic characterization demonstrates intrinsic disorder in the N-terminus and linker regions

To experimentally determine the unstructured content of the protein, we used CD and NMR to assess the secondary structure. Our CD secondary structure assignment for ovalbumin were similar to the published x-ray crystal structure under our conditions (54). In Cdt1, spectral features indicate a large portion of the protein is neither helical nor sheet, and approximately 50% of the protein is in the “random coil” region (55). We also performed ^1^H/^15^N HSQC NMR to validate our CD results. The number of peaks, their broadness and their location similarly demonstrate a high disorder content (56). This data confirms all but the WH domains are disordered in human Cdt1. Microtubule-binding proteins like Tau, Knl1(57), and TPX2(58), are well-reported to contain intrinsically disordered regions as well, all consistent with scaffold functions. Intrinsic disorder can also facilitate engagement of the proteasome (59), which in the case of Cdt1, could again aid in rapid degradation throughout the cell cycle.

### Hydrodynamic and scattering experiments show Cdt1 is a monomeric, mixed-folded protein with significant conformational heterogeneity

After validating Cdt1 contains disordered regions at the N-terminus and in between the WH domains, we next used scattering and chromatography techniques to further investigate biophysical properties. We used Dynamic Light Scattering (DLS) to estimate the radius of hydration (R(h)) as well as polydispersity of Cdt1 in solution. Ovalbumin showed similar R(h) values in our system as with published results (60). For Cdt1, DLS demonstrated the protein was 99% monomeric, had a calculated R(h) of 3.7 nm, and was 40% polydisperse (**Figure 3A**). This relatively high level of polydispersity is consistent with a mixed-folded protein with many different conformations and also is consistent with thermally instability at relatively low temperatures (61). In a set of 124 protein-protein complexes, one group proposed that increased conformational heterogeneity increased interface “recognition” to promote complex formation (62), which could conceivably also apply to Cdt1.

**Figure 3.**
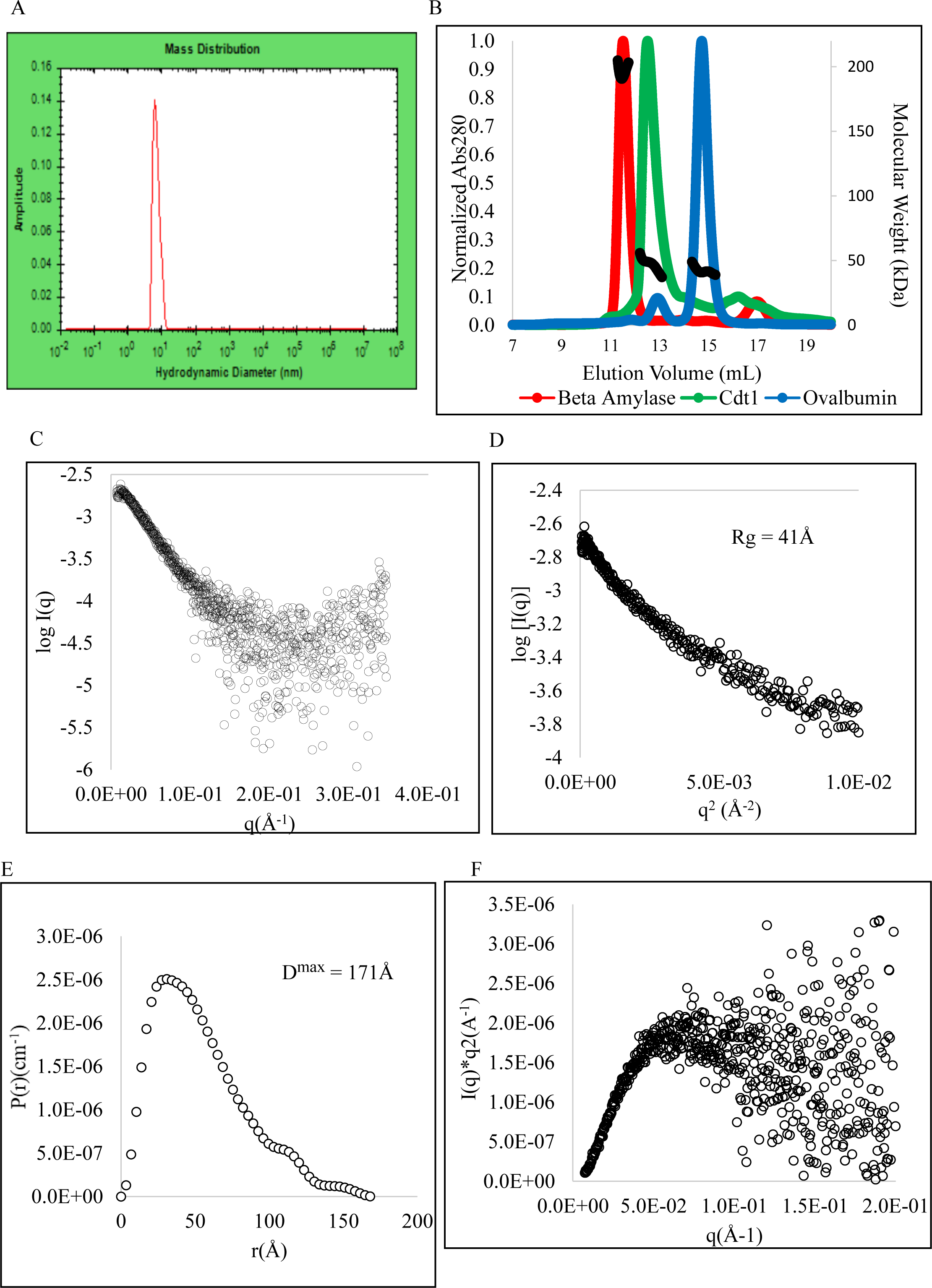
Hydrodynamic and scattering analyses of Cdt1. (A) DLS mass distribution for Cdt1. Amplitude is plotted as a function of the log of hydrodynamic diameter (nm). (B) SEC-MALS chromatogram of Cdt1 and model proteins. Normalized concentration (Abs280) and calculated MW (kDa) are shown as a function of elution volume (mL). Red is beta amylase, green is Cdt1, and blue is ovalbumin. (C) Normalized scattering plot of Cdt1. Log scattering intensity [Log(I)] is plotted as a function of scattering angle (Å^−1^). (D) Guinier plot of Cdt1. Natural log of intensity (Ln(I) in relative units) is plotted as a function of scattering angle (Å^−1^). (E) Distance distribution function. The distribution of distances between all pairs of points within the protein (P(r)) is plotted as a function of radius (Å). (F) Kratky plot. I*q^2^ is plotted as a function of scattering angle (q).

Size exclusion chromatography coupled with multi-angle light scattering (SEC-MALS) can orthogonally determine the radius of hydration (R(h)), molecular weight, therefore, oligomeric state (63). Our SEC system demonstrated an excellent correlation between our experimental elution volume and previously published R(h) values for SEC standard/model proteins (36,60). Analysis of six protein standards demonstrated our system was accurate to, on average, ±10% in mass from published values (**Figure S2B, Table S3**). When we tested Cdt1, it eluted significantly earlier than would be expected for a well-folded protein of similar molecular weight like ovalbumin. However, it eluted significantly after the void volume and the tetrameric beta amylase, consistent with a partially folded protein rather than aggregate. Assuming no column resin interactions, based on interpolation of the SEC standards (64), we calculated the R(h) of Cdt1 to be 4.4 nm (**Table 1, Figure S2C-F**). MALS determined this Cdt1 construct to be 46 kDa, within 10% of the theoretical monomer MW (**Figure 3B**). Because the calculated molecular weight remained constant throughout the peak and therefore, at a range of concentrations, we infer there may be limited inter-molecular self-association or transient oligomers under our conditions.

To gain further build on the DLS and SEC-MALS experiments, we performed size exclusion chromatography coupled with small-angle x-ray scattering (SEC-SAXS, (60)). Using SEC upstream of SAXS allows for improved buffer subtraction, removal of aggregates, and separation of oligomers which could otherwise confound analysis (65,66). The SAXS results for ovalbumin were comparable to previously published values (**Figure S3A-D**, (67)). Normalized scattering is shown in **Figure 3C**. For Cdt1, under our conditions, analysis of the Guinier region (**Figure S4A** & **4B**) shows an Rg of 41 Å and a deviation from linearity (**Figure 3D**), as is expected for a partially disordered protein with a q_max_*R_g_ > 1.1(68,69). The R_g_ and D_max_ values (**Figure 3D** & **3E**) were also noticeably larger than globular ovalbumin (**Table 1, Table S4**). For Cdt1, the P(R) and Kratky plots (**Figure 3E** & **3F**) are both consistent with a partially unfolded protein. These solution scattering results, in addition to the Rg/R(h) ratio, validate Cdt1 contains both significant folded and unfolded regions. The calculated Rg values were also independent of protein concentration across the SEC elution peak (**Figure S4C**). In agreement with the MALS calculations, our SAX data is consistent with the interpretation that there was no significant inter-molecular interactions or self-association observed under the conditions tested, which otherwise would have shown a change in MW or Rg as a function of elution time. Both our SEC-MALS and SEC-SAXS data are interpreted as Cdt1 being monomeric with minimal inter-molecular self-interactions, even at concentrations as high as 110 µM (see Methods) *in vitro*.

Cdt1’s most well-characterized mitotic binding partners, including the Ndc80 and Ska1 protein complexes, have been shown to homo-oligomerize and function around microtubules or in the presence of other binding partners (70,71). Ndc80 can homo-oligomerize on its own, consistent with its load-bearing role at microtubules. Our data shows, in contrast, that *in vitro* Cdt1 cannot independently form higher order oligomers in the absence of other kinetochore components. And at the preRC complex, a monomeric state would likely be required to ensure only one licensing step. Both functions imply binding partner association, not high local concentration in isolation, drive Cdt1 complex formation at Ndc80, consistent with its function as a scaffold protein.

### Rigid body modeling suggests the two folded domains of Cdt1 are spatially close to each other in “closed” conformations

To better understand the structural conformations of folded and unfolded regions together, we proceeded by combining experimental and computational approaches using SAXS/MD/MC from the web server SASSIE (72,73). By combining experimental SAXS data with molecular simulations, “conformational space” can be better sampled and SAXS/MD/MC can access structural information otherwise difficult to capture solely by experimental means (74). The SASSIE workflow is shown in **Figure S5A** and additional simulation details are in the Methods section. In brief, using our input experimental SEC-SAXS data with SASSIE, we 1) created an initial model using the WH domains as rigid bodies, 2) generated an ensemble of structures, 3) calculated their theoretical SAXS curves, and then 4) analyzed the best fit conformations using Rg and χ^2^ filters. With this approach, we aim to understand possible conformations for mixed-folded systems like Cdt1. Using our experimental secondary structure content analysis, we defined the N-terminal and linker regions as entirely flexible. We first generated a total of 250,000 structures, of which 13,724 were accepted based on steric hinderance. This “conformational space” is shown as black dots in **Figure 4A** and black mesh in **Figure 4B**. We then filtered the data according to models that best fit both our experimental Rg and χ^2^ fit to the scattering curve. Only 3,945 of those had a χ^2^ < 3 to the experimental scattering data, shown in blue in **Figure 4A** and **4B**.

**Figure 4.**
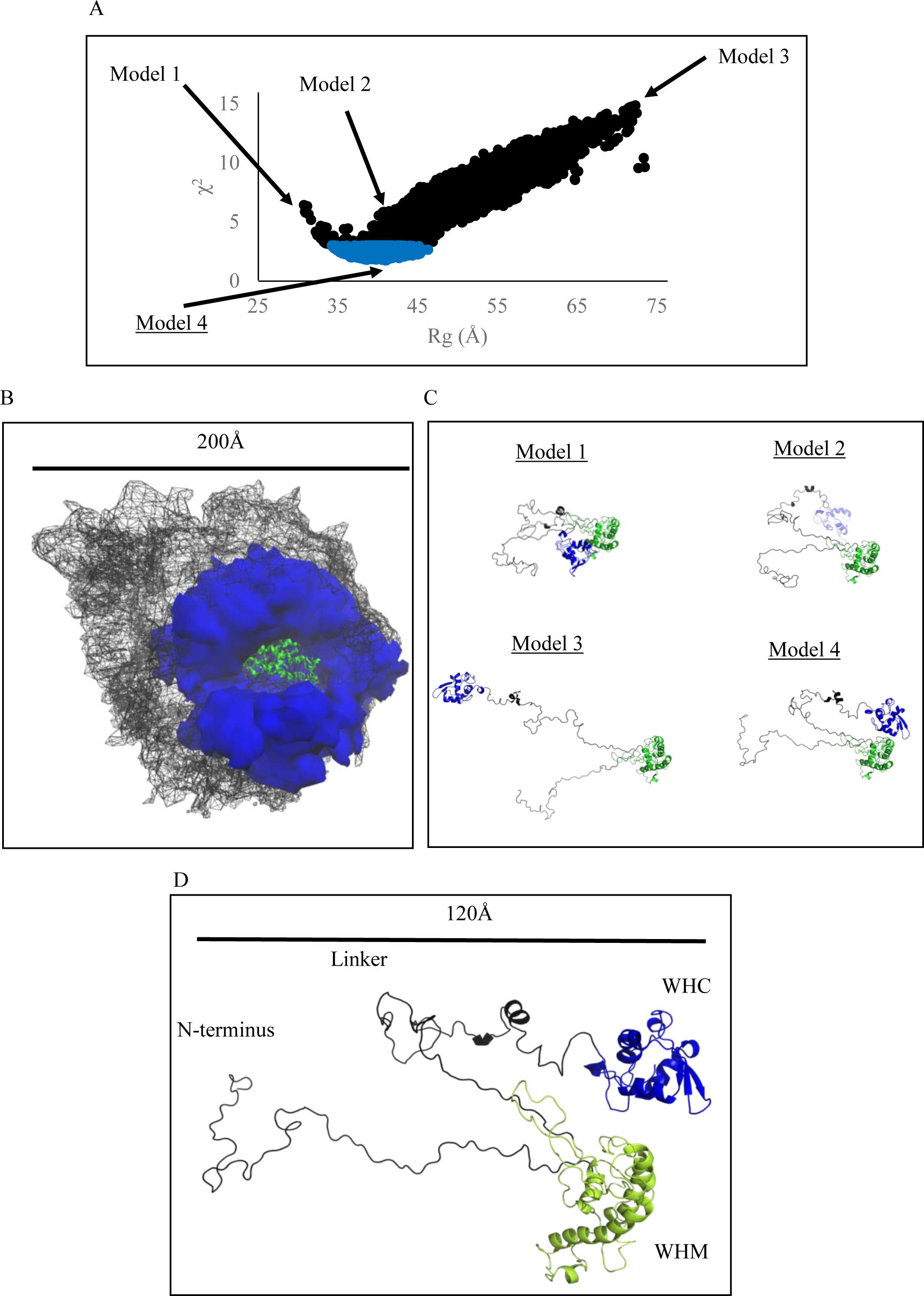
SAXS/SASSIE rigid body modeling of Cdt1. (A) Plot of Rg vs. χ^2^. All generated models are shown in black where models with a χ^2^ value less than 3 are colored blue. Representative Models are indicated by arrows and labels. (B) Ensemble of structures. Total accepted steric conformational space is shown in black mesh while experimental well-fit conformations are shown in blue. (C) Representative conformations. The WHM domain is fixed and shown in green. The WHC domain is in blue. N-terminal, and linker regions are in black. Models are elaborated in text. (D) Enlargements of a representative best fit Model 4 from panel C.

Representative models are shown in **Figure 4C** and **Table 2**. Best, worst, and average computational fits to experimental data are shown in **Figure S5B. Model 1**, with an extremely compact conformation, had an Rg that was too low to fit our SAXS data. **Model 2**, while having an accurate Rg, did not have a low χ^2^ fit to our q vs. I(q) plot. Similarly, the **Model 3** “extended” conformations, with the two domains being spaced approximately 140 Å apart, was among the worst fits to our data. This contrasts with proteins like Ataxin-3, where similar SAXS/MD analyses (75,76) showed the domains and disordered tail are all separated far apart from each other. Because Ataxin-3 functions as a deubiquitinase, these conformations are thought to help bind and cleave across extended poly-ubiquitin chains.

**Table 2:** Rg, χ^2^, and WHD center-of-mass distances of representative model structures.

| Model | R<sub>g</sub> (Å) | $\chi^2$ | Center-of-mass distance between WH domains (Å) |
| --- | --- | --- | --- |
| 1 | 31 | 6.3 | 24 |
| 2 | 40 | 5.5 | 59 |
| 3 | 72 | 14.8 | 153 |
| 4 | 41 | 1.7 | 39 |

The best fit data, **Model 4**, is shown in blue in **Figure 4B** and **4C** and has an Rg of 41 Å, in agreement with our SEC-SAXS data. Conformations like Model 4 are those where the WHM and WHC domains are close to each other, with the N-terminal and linker regions being extended into solution (**Figure 4D**). A comparison of the folded Cdt1 domains with the ovalbumin shows that it is likely these linker and N-termini regions are likely responsible for the increased R(h) and Rg (**Figure S5C**). Despite having stretched of 67 and 54 amino acid disordered regions respectively, and no signs of inter-molecular self-interaction via scattering techniques, the two WH domains are only approximately 20Å apart from each other at their closest in **Model 4**. We refer to this best-fit conformation as “folded over”.

We also calculated the inter-domain distances of this ensemble from their center-of-masses. The WHM and WHC boundaries used in **Figure 1A**, and used historically, were used as domains. Because it is probable that each WH domain binds to distinct sites in its protein complexes, studying Cdt1’s inter-domain center-of-mass would validate our previous simulations and be useful for future comparisons. Calculations for each of the four models are shown in **Table 2**. The distribution of inter-domain distances was broad, given the range and number of structures derived from the Monte Carlo simulations (**Figure 5A**). Only structures with inter-domain distances between 35-52Å had χ^2^ <2 (**Figure 5B**). This distribution was slightly larger for those structures with χ^2^ <3 (**Figure 5C**), as expected. And for χ^2^>5, only poor-fitting structures were abundant (**Figure 5D**). From this analysis, χ^2^ best fit values, and χ^2^ worst fit values both agree with our SASSIE-generated ensemble, and consistent with “folded over” conformations as a well-fit structures. This center-of-mass analysis may aid in future intra-molecular FRET probe studies. Unfortunately, yeast Cdt1 has an additional folded domain (77) and is only 39% similar to human Cdt1, making structural comparisons to those at the preRC difficult (2).

**Figure 5.**
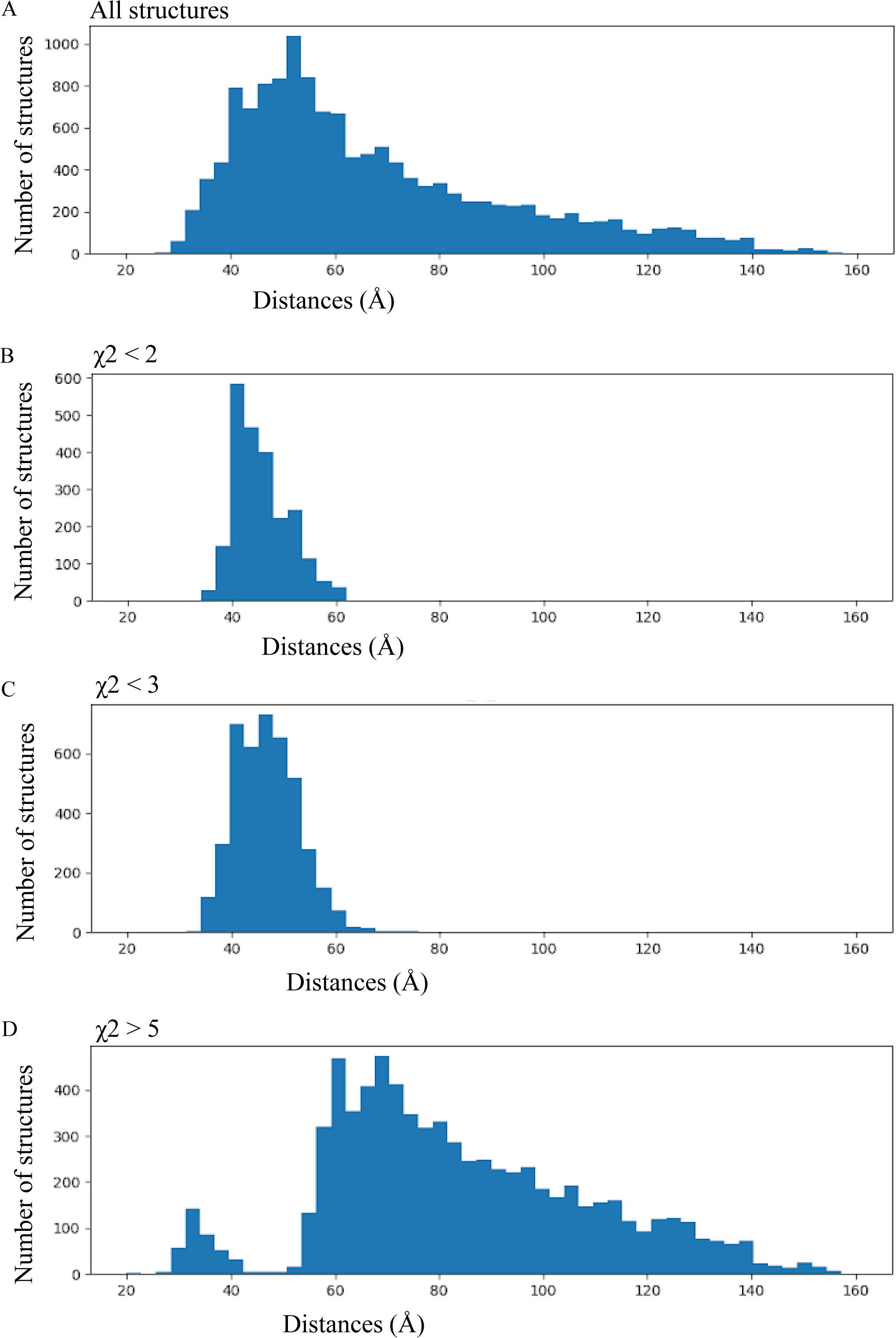
WHM/WHC center-of-mass, inter-domain histograms of Cdt1. (A) Number of structures in our ensemble as a function of inter-WHD center-of-mass distances. (B) Number of structures in our ensemble as a function of inter-domain center-of-mass distances with χ^2^ less than 2. (C) Number of structures in our ensemble as a function of inter-domain center-of-mass distances with χ^2^ less than 3. (D) Number of structures in our ensemble as a function of inter-domain center-of-mass distances with χ^2^ greater than 5.

These” folded over” conformations could possibly exhibit some level of steric hinderance such that it could be less likely to bridge kinetochore components without some level of “opening up”. In SAXS/MD/MC studies of TraI, the domains and linkers adopt similar “folded over” conformations, with the folded domains close to each other and the linker extended out in solution. TraI functions as a helicase and scaffold protein for the DNA strand transfer machinery during conjugation n bacteria (78). Based on their data, the authors suggested this inter-domain proximity could be responsible for the observed negative cooperation between the nickase and ssDNA binding domains, which otherwise would extend apart. (79). Multi-domain proteins that fold over onto themselves can sometimes be known to have auto-inhibitory functions (80,81). Based on our SAXS/MD/MC data, and in the context of other data on mixed-folded proteins, we hypothesize that Cdt1 could potentially exist in solution in some form of an auto-inhibitory state. It appears that Cdt1 may have to undergo a relatively large conformational change to extend and bridge multiple components of the kinetochore. This work further enables future reductionist approaches to study complex formation with Ska/Ndc80, and MCM, and also other mixed-folded protein systems.

## Author Contributions

K.P.S. and D.V. initiated the project. K.P.S. managed the project and designed the experiments. K.P.S., A.R., and K.M.V. expressed and purified proteins. M.T. collected and analyzed NMR data. M.C. collected and analyzed TIRF data. M.G.P. collected and analyzed DSF data. K.M.V. collected and analyzed DLS data. S.C. collected SAXS data. A.A.G. assisted in CD and SEC-MALS data collection and analysis. K.P.S. and J.E.C. performed simulations, analysis, and interpretation. D.V., A.A.G., and J.E.C. provided additional intellectual input. K.P.S. wrote the manuscript with assistance from D.V. and input from all authors.

## Acknowledgements

We thank Dr. Cara Gottardi for her support and mentorship. We thank Anita Nair-Varma for help with laboratory operations. We thank Dr. Sergii Pshenychnyi of the Recombinant Protein Production Core at Northwestern for supplying TEV protease. We thank Dr. Ronald Soriano, James Casey, and the Northwestern Keck Biophysics Facility for assistance with data collection. This work used resources of the Northwestern University Keck Biophysics Facility, supported by NCI-CCSG-P30-CA060553 awarded to the Robert H. Lurie Comprehensive Cancer Center. Use of the Advanced Photon Source was supported by the US Department of Energy, Office of Science, Office of Basic Energy Sciences, under Contract No. DE-AC02-06CH11357. Use of the Pilatus 3 1M detector was provided by grant 1S10OD018090-01 from NIGMS. This work was supported by NIGMS grant R01GM135391 to D.V.

## Supporting Material

**Figure S1.**
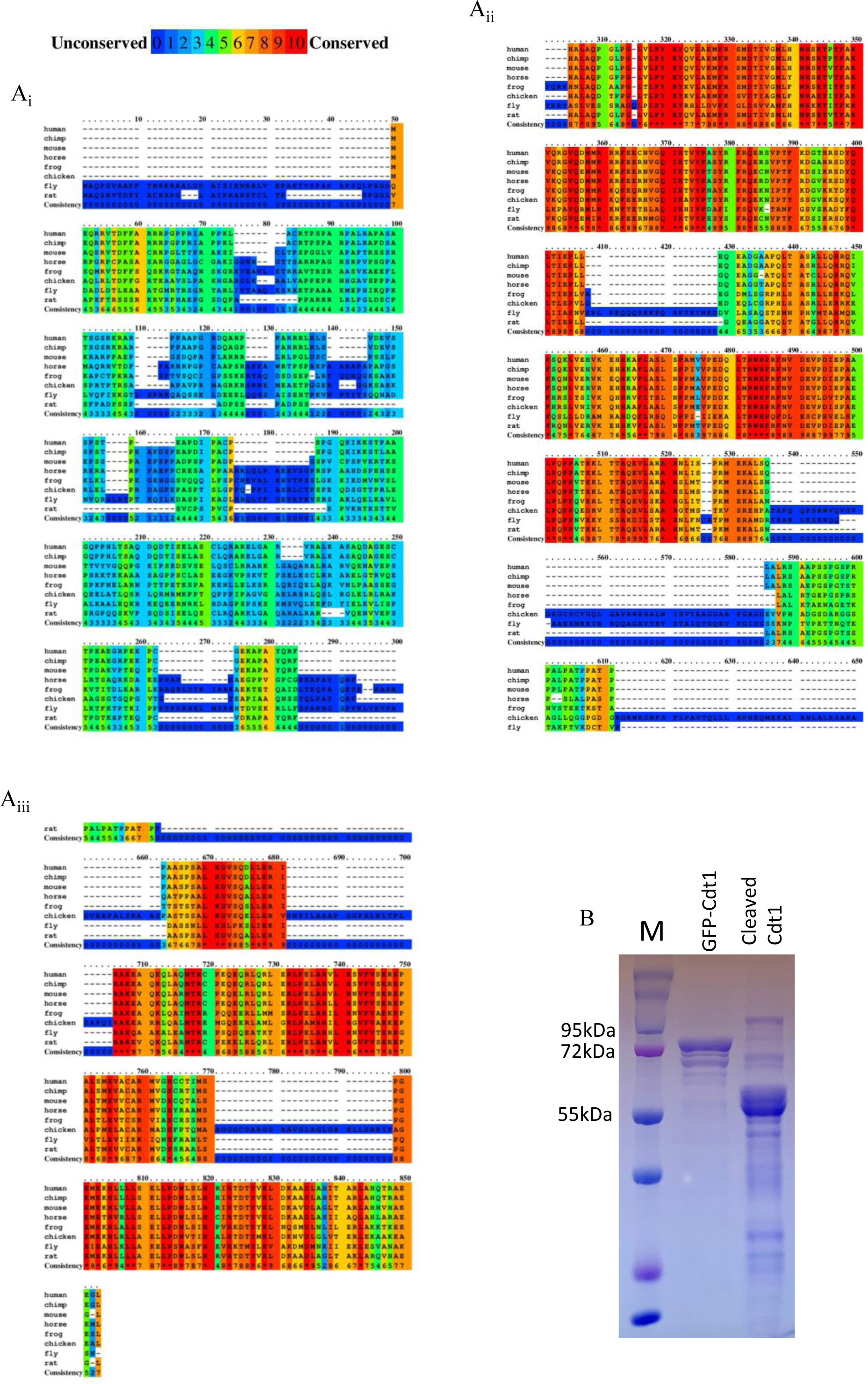
Cdt1 amino acid sequence analysis and SDS-PAGE of Cdt1 constructs. (A) Amino acid sequence alignment of Cdt1. Human, chimpanzee, mouse, horse, frog, chicken, fly and rat sequences are aligned top to bottom. Conservation is colored blue for least conserved and red for most conserved. (B) SDS-PAGE of one production of Cdt1. Lanes are 1) MW ladder, 2) GFP-Cdt1, 3) cleaved Cdt1.

**Figure S2.**
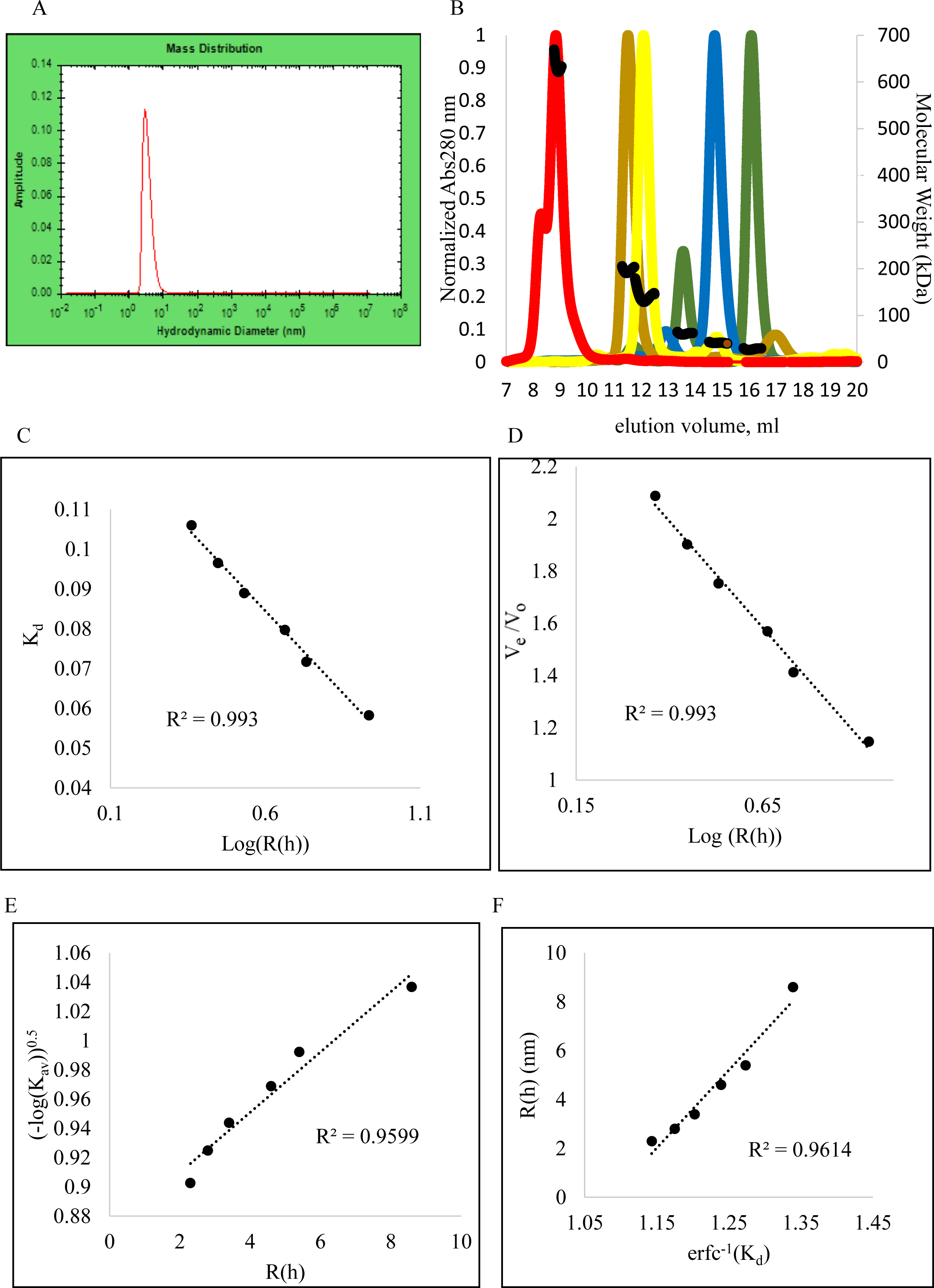
DLS and SEC-MALS of protein standards. (A) DLS mass distribution profile of ovalbumin. Amplitude is plotted as a function of the log of hydrodynamic diameter (nm). (B) SEC-MALS chromatogram for protein standards. Normalized concentration (Abs280) and calculated MW (kDa) are shown as a function of elution volume (mL). (C) A plot of K_d_ as a function of (LogR(h)). (D) A plot of V_e_/V_o_ as a function of Log(R(h)). (E) A plot of (-log(K_av_))^1/2^ as a function of R(h). (F) A plot of R(h) as a function of _erfc −1(Kd)._

**Figure S3.**
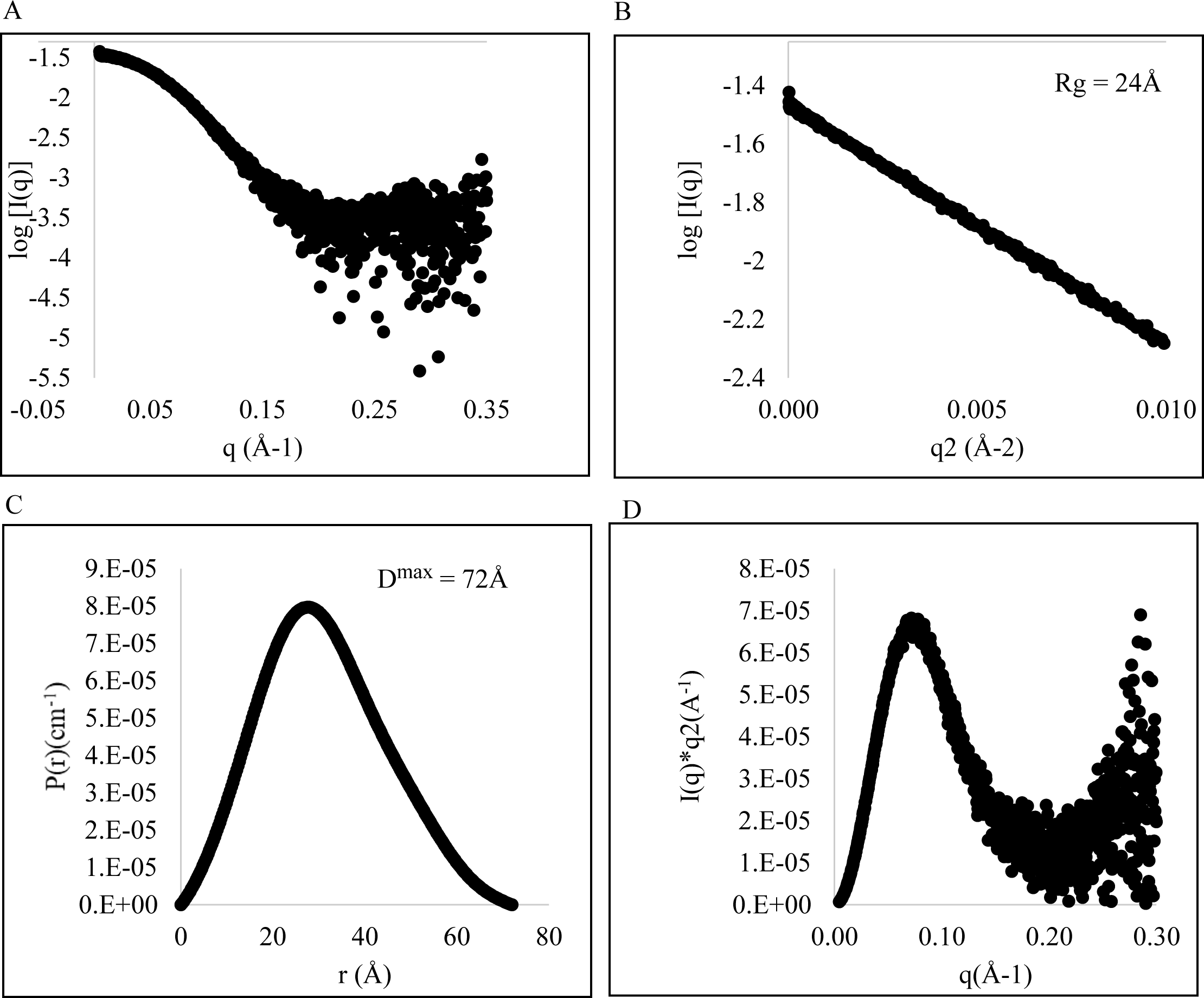
SEC-SAXS of ovalbumin. (A) Normalized scattering plot of ovalbumin. Log scattering intensity (Log(I) in relative units) is plotted as a function of scattering angle (Å^−1^). (B) Guinier plot of ovalbumin. Natural log of intensity (Ln(I)) is plotted as a function of scattering angle (Å^−1^). (C) Paired distance distribution function. Distribution of distances between all pairs of points within the protein (P(r) in relative units) is plotted as a function of radius (Å). (D) Kratky plot of ovalbumin. I*q^2^ is plotted as a function of scattering angle, q.

**Figure S4.**
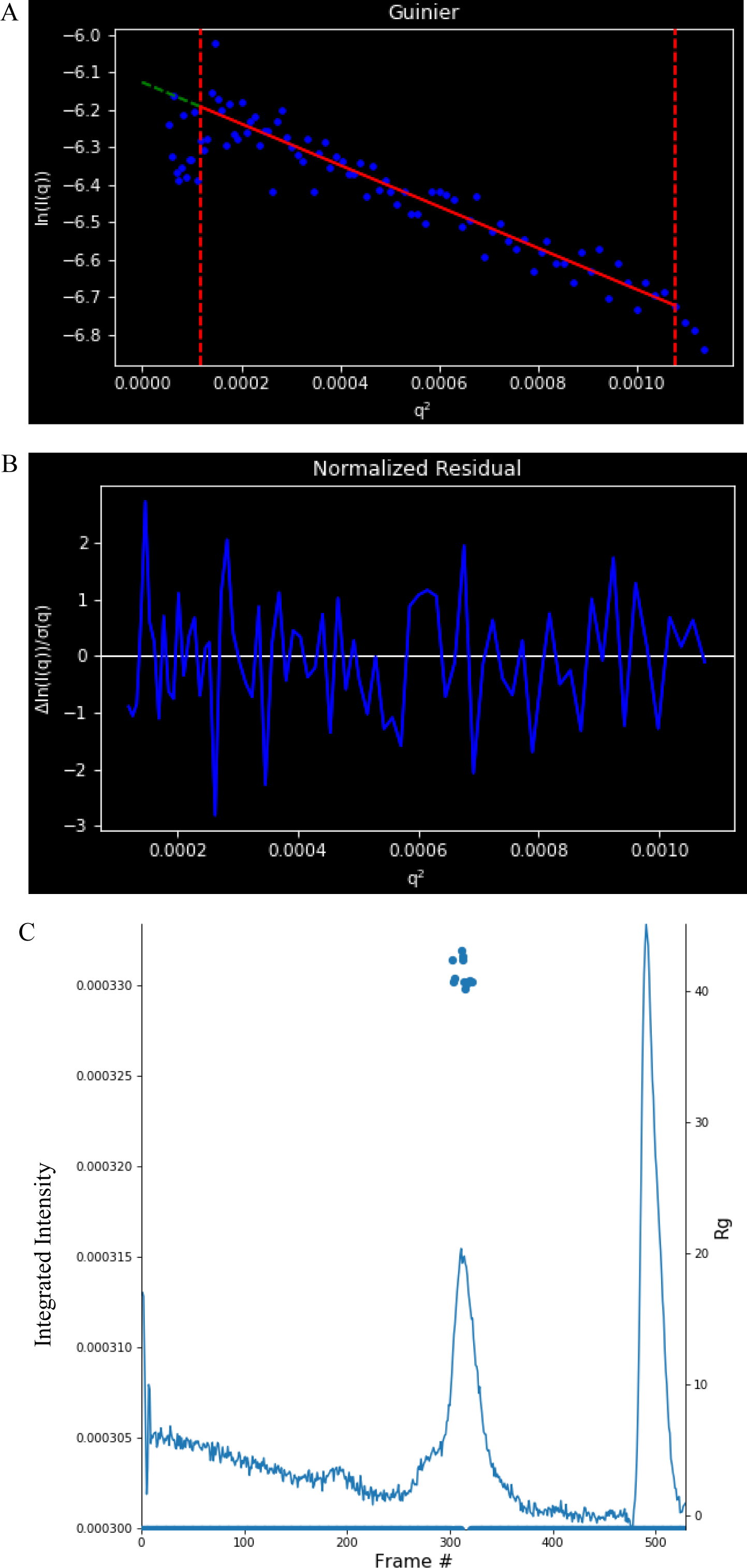
Guinier analyses of Cdt1. (A) Guiner plot. Ln(I(q)) is plotted as a function of q^2^. Boundaries are shown as red dashed lines. Linear fit of data shown in solid red line. (B) Normalized residual. Δln(I(q))/σ(q) is plotted as a function of q^2^. (C) Rg across the SEC elution peak for Cdt1. Integrated intensity and Rg are plotted as a function of Frame #.

**Figure S5:**
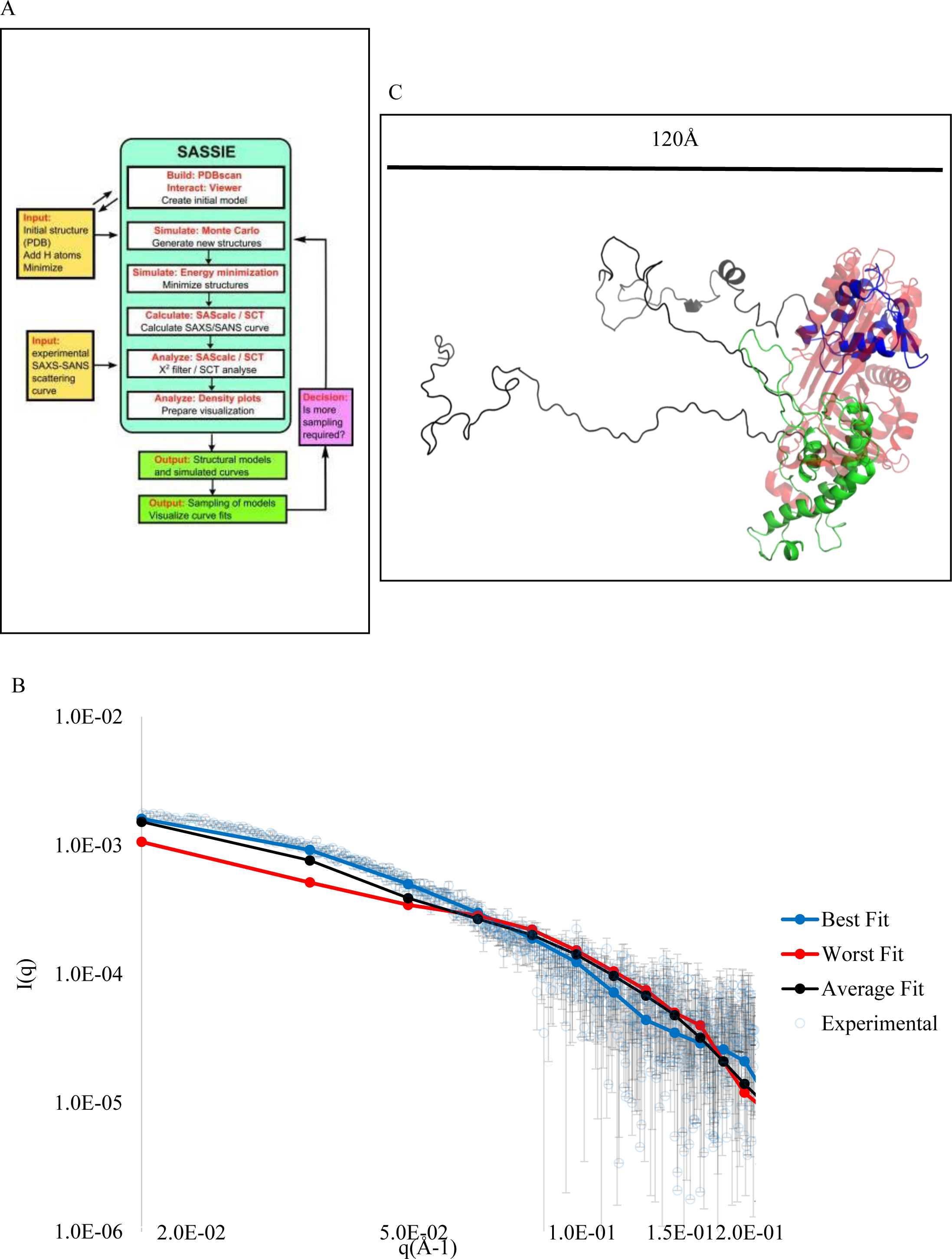
SASSIE workflow and SASSIE model fits to Cdt1 scattering data. (A) SASSIE workflow, taken from (73). (B) Overlay of Cdt1 Model 4 and ovalbumin. Cdt1 WHM domain is in green, the WHC domain is in red, the N-terminal and linker regions are in black, and ovalbumin is shown as translucent red. Scale bar is 120 Å. (C) Raw scattering data and fits to models. Experimental data is in grey, worst fit is in red, average fit is in black, and best fit is in blue.

**Table S1:** Ovalbumin Thermal Denaturation.

| Method | Melting Temperature (°C) |
| --- | --- |
| CD (222 nm) | 77 |
| FL (Intrinsic Fluorescence) | 76 |

**Table S2:** Ovalbumin Secondary Structure Content.

| Method | α-Helix (%) | β-Sheet (%) | Other (%) |
| --- | --- | --- | --- |
| CD | 23 | 21 | 56 |
| X-Ray Structure (54) | 27 | 34 | 39 |

**Table S3:** SEC-MALS standards.

| Sample | Elution Volume (mL) | Theoretical Molecular Weight (kDa) | MALS Calculated Molecular Weight (kDa) | Published R(h) (nm) |
| --- | --- | --- | --- | --- |
| Ethanol | 19.51 | 0.05 | N/A | N/A |
| Carbonic Anhydrase | 16.08 | 30 | 28 | 2.3 (60) |
| Ovalbumin | 14.65 | 43 | 51 | 2.8 (60) |
| Bovine Serum Albumin | 13.50 | 67 | 62 | 3.4 (82) |
| Cdt1 | 12.51 | 51 | 46 | 4.4* |
| Alcohol Dehydrogenase | 12.09 | 150 | 138 | 4.6 (83) |
| Beta Amylase | 10.88 | 224 | 198 | 5.4 (84) |
| Thyroglobulin | 8.84 | 669 | 674 | 8.6 (85) |
| Blue Dextran | 7.70 | 2000 | N/A | N/A |

N/A=Not Applicable. (\*) denotes interpolation from this study.

**Table S4:** Biophysical Properties of Ovalbumin.

| Parameter | Value |
| --- | --- |
| DLS R(h) (Å) | 26 |
| SEC Aggregate R(h) (Å) | 28 |
| SAXS R<sub>g</sub> (Å) | 36 |
| SAXS D<sub>max</sub> (Å) | 72 |
| MALS Calculated MW (kDa) | 42 |
| Theoretical MW (kDa) | 43 |

